# Expanding the depth and sensitivity of cross-link identification by differential ion mobility using FAIMS

**DOI:** 10.1101/2020.04.23.057091

**Authors:** Lennart Schnirch, Michal Nadler-Holly, Siang-Wun Siao, Christian K. Frese, Rosa Viner, Fan Liu

**Author notes:** Correspondence (F. L.).

## Abstract

In cross-linking mass spectrometry, the depth and sensitivity is often limited by the low abundance of cross-links compared to non-cross-linked peptides in the digestion mixture. To improve the identification efficiency of low abundant cross-links, here we present a gas-phase separation strategy using high field asymmetric waveform ion mobility spectrometry (FAIMS) coupled to the Orbitrap Tribrid mass spectrometers. By enabling an additional peptide separation step in gas phase using the FAIMS device, we increase the number of cross-link identification by 23% for a medium complex sample and 56% for strong cation exchange-fractionated HEK293 cell lysate. Furthermore, we show that for medium complex samples, FAIMS enables the collection of single-shot cross-linking data with comparable depth to the corresponding sample fractionated by chromatography-based approaches. Altogether, we demonstrate FAIMS is highly beneficial for XL-MS studies by expanding the proteome coverage of cross-links while improving the efficiency and confidence of cross-link identification.

## Introduction

Proteins do not act alone. They interact with each other by forming stable protein complexes and dynamic signaling cascades through protein-protein interactions.^1,2^ Probing the interaction interfaces of native protein complexes and profiling large-scale protein-protein interaction networks are of great importance to gain molecular understandings of biological processes. Chemical cross-linking combined with mass spectrometry (XL-MS) emerged as a powerful technique to tackle these questions.^3,4^ In XL-MS, cross-linking reagents are applied to proteins and complexes to covalently link the residue pairs in close proximity in the three-dimensional space. Subsequently, cross-linked proteins are proteolytically digested into a mixture of peptides that are primarily composed of non-cross-linked linear peptides, mono-links, loop-links and cross-links.^3,5,6^ Among all different types of peptides, cross-links are of the most importance as they render a maximum distance constraint between the two linked residues (i.e., the maximum distance given by the length of the cross-linker spacer arm).^7^ This distance information is essential to obtain structural information of proteins and protein complexes as well as to build protein connectivity networks.^8,9^

Despite the significant importance, cross-links are often lower abundant compared to other types of peptides in the digestion mixture, hindering efficient cross-link identification directly from the total peptide population.^6^ Therefore, a crucial step in XL-MS sample preparation is cross-link enrichment prior to liquid chromatography couple to mass spectrometry (LC/MS) analysis.^5,10^ Current technologies to enrich cross-linked peptides include chromatography-based peptide separation strategies such as size exclusion chromatography (SEC)^11^ and strong cation exchange chromatography (SCX)^8^, as well as affinity-based enrichment making use of affinity-tagged cross-linkers^12^. These fractionation and enrichment strategies are essential to XL-MS studies, especially for samples of medium and high complexity (i.e., macromolecular assemblies and proteome samples).

In addition to in-solution-based separation, peptides can also be fractionated in the gas phase using ion mobility spectrometry which has been proven to be orthogonal to online LC separation.^13–15^ Differential ion mobility separates ions depending on their physiochemical properties such as mass, shape, center of mass, dipole moment and charge.^16,17^ Recently, a new generation ion mobility device was released, enabling high-field asymmetric waveform ion mobility spectrometry (FAIMS) with improved characteristics compared to previous generations.^18^ The device is a front-end instrument coupled to Orbitrap mass spectrometers (FAIMS Pro™ interface from Thermo Fisher Scientific). In FAIMS, ions are transferred by the carrier gas (N2), while passing through an alternating electric field between the two electrodes. Ions that show a different mobility in high and low electric fields are displaced towards one of the electrodes. By hitting the electrode, these ions are hindered from passing through the device. To select ions for analysis, an additional direct current (compensation voltage, CV) is applied. This adjustable CV is able to partially compensate for the displacement of certain ions and enables them to leave the device and enter the mass spectrometer. Applying different CVs either in successive MS runs (external CV stepping) or within one MS run (internal CV stepping) allow ions to be analyzed in subsets. Therefore, CV is the most important parameter to fine-tune for different proteomic experiments.^19–21^

Because of the recent technical advancements, the new generation of FAIMS exhibits high ion transmission (>70% of ions transmitted at optimal CV), prolonged spray stability (120 hours of continuous use), as well as robustness to different setups (column diameter and flow rate).^18^ It has been demonstrated that FAIMS improves the depth and sensitivity of proteome coverage as well as the accuracy of protein quantification in quantitative proteomics. These advances are benefited from increased sensitivity, reduction in co-eluting or fragmenting peptides and suppression of interfering +1 charged features.^18,19^

Based on these previous achievements, we reason that FAIMS may be of great value in increasing sensitivity also for cross-linked peptides. To assess its performance in detail, here we applied FAIMS to XL-MS studies of protein samples of different complexity. Our data show that FAIMS significantly increases the identification of cross-linked peptides from unfractionated medium complex samples by more than 20%. Furthermore, FAIMS improves cross-link identification to a similar level as if SEC is used, thus offering an easy alternative to chromatography-based fractionation for medium complex samples. For high complex samples, as illustrated by SCX fractionated cross-linked cell lysates, FAIMS increases the number of cross-linked peptides and protein pairs by up to two-fold in early fractions and approximately 40% in later fractions. Finally, based on extensive testing of FAIMS settings, we suggest optimal sets of CVs for samples in various complexity.

## Materials and Methods

### Sample preparation and cross-linking

To generate the eight cross-linked protein mixture, horse myoglobin, bovine hemoglobin, bovine serum albumin, bovine beta-casein, bovine beta-lactoglobulin, bovine cytochrome-c, chicken ovotransferrin and chicken ovalbumin (all purchased from Sigma Aldrich, except the last two which were purchased from GE healthcare as part of the Gel Filtration High Molecular Weight Kit) were prepared individually as 1 mg/ml solutions in 20 mM HEPES pH 7.8 prior to cross-linking. HEK293T cells were lysed in 20 mM HEPES, 150 mM NaCl, 1.5 mM MgCl2, 0.5 mM dithiothreitol (DTT) pH 7.8 in a 5 ml Dounce homogenizer (20 passes of a tight-fitting glass piston at 1000 rpm). Cell lysate was cleared by centrifugation at 1000 x g for 5 min at 4 °C. Protein concentration was estimated at 1 mg/ml based on cell counts prior to lysis.

Chemical cross-linking was performed with 0.5 mM disuccinimidyl sulfoxide (DSSO) (Thermo Fisher Scientific, resuspended freshly in anhydrous DMSO to 50 mM) and incubated at room temperature for 1 h. The reaction was stopped by addition of 50 mM Tris, pH 8.0 final for 30 min.

Cross-linked samples were denatured using 8M Urea, reduced with DTT (5 mM, 1 h at 37 °C) and alkylated with chloroacetamide (CAA, 40 mM, 30 min at room temperature in the dark). Proteins were sequentially proteolyzed with Lys-C (1:75 w/w, 2h at 37 °C) and trypsin (1:100 w/w, overnight at 37 °C). Digests were desalted using Sep-Pak C8 cartridges (Waters), dried under vacuum, and reconstituted either in 0.05% (v/v) trifluoroacetic acid (TFA), 1% (v/v) acetonitrile (ACN) for subsequent LC/MS measurements or 10% (v/v) formic acid (FA) prior to fractionation by strong cation exchange (SCX) chromatography. Peptide concentrations were determined using Pierce Quantitative Colorimetric Peptide Assay (Thermo Fisher Scientific). For the eight protein mixture, tryptic peptides of respective eight proteins were mixed 1:1 to obtain a mixture that resembles a medium complex sample.

### Strong cation exchange (SCX) fractionation of cross-linked peptides

Fractionation was performed on an Agilent 1260 Infinity II UPLC system equipped with a PolySULFOETHYL-A™ column (100 x 4.6 mm, 3μm particles, PolyLC inc.). 2 mg HEK293T cell digest was loaded. Solvent A consisted of 0.05% FA in 20% ACN, and solvent B consisted of the same buffer supplemented with 0.5 M NaCl. The SCX gradient was as follows: 0–0.01 min (0–2% B); 0.01–8.01 min (2–3% B); 8.01–14.01 min (3–8% B); 14.01–28 min (8–20% B); 28–48 min (20–40% B); 48–68 min (40–90% B); 68–74 min (90% B); and 74–95 min (0% B). Fractions were collected every 30 seconds starting from minute 35. Six equally spaced late SCX fractions along the gradient were selected for LC/MS experiments.

### Size exclusion chromatography (SEC) fractionation of cross-linked peptides

Size exclusion chromatography was performed on a quaternary Agilent 1290 Infinity II UPLC system equipped with a SEC column (Superdex Peptide 3.2/300, GE healthcare). 40 μg of the eight protein mixture was loaded and separated at a flow rate of 0.05 ml/min with running buffer (30%ACN, 0.1%TFA). The early five (20-30 min), cross-link enriched fractions were selected for LC/MS experiments.

### Liquid Chromatography and Mass Spectrometry

Samples were separated by RP-HPLC using a Thermo Scientific™ Dionex™ UltiMate™ 3000 system connected to a PepMap C-18 trap-column (0.075 mm x 50 mm, 3 μm particle size, 100 Å pore size (Thermo Fischer Scientific)) and an in-house packed C18 column for reversed phase separation (column material: Poroshell 120 EC-C18, 2.7 μm (Agilent Technologies)) at 300 nL/min flow rate. The crosslinked samples were analyzed on the Orbitrap Fusion Lumos mass spectrometer with or without FAIMS Pro™ device (Thermo Fisher Scientific) with Instrument Control Software version 3.3. Reverse phase separation was accomplished using a 60, 90, 120 min separation gradient (plus 30 min equilibration phase) for the eight protein mixture or 180 min gradient (separation for 160 min, plus 20 min equilibration phase at the end) for HEK293T cell lysate. Separation gradient: 4-40% solvent B (A: water, 0.1% FA; B: 100% ACN, 0.1% FA).

Cross-linked samples were analyzed using stepped collision energy (SCE) HCD-MS2-MS3 acquisition strategy. MS2 sequencing events were triggered for precursors at charge states +4 to +8 and mass-difference-dependent CID-MS3 acquisitions were triggered if a unique mass difference (Δ=31.9721) was observed in the (SCE) HCD-MS2 spectrum for two of the signature peaks. MS1 and MS2 scans were acquired in the Orbitrap with a respective mass resolution of 120,000 and 60,000 whereas MS3 scans were acquired in the ion trap. For MS3 scans, precursor isolation windows were set to 1.6 m/z at MS1 level and 2.5 m/z at MS2 level. The normalized collision energy was set to 21-27-33% for (SCE) HCD-MS2 scans and 35% for CID-MS3 scans. Data were acquired using top speed of 5 sec for non FAIMS or single CV experiments and 2 sec/CV for internal stepping FAIMS experiments. Dynamic exclusion was set to 30, 45 or 60 sec. for increasing gradient length. For protein identification and abundance estimation, single-shot LC-MS experiments of unfractionated cross-linked HEK293T cell lysate were performed using an LTQ Orbitrap Elite mass spectrometer (Thermo Fisher Scientific) and 180 min linear gradients as described above.

### Data analysis

Spectral raw data files were analyzed using Proteome Discoverer 2.4 software (Thermo Fisher Scientific) with XlinkX node 2.0 for cross-linked peptides and SEQUEST HT search engine for unmodified peptides and mono-links. MS1 ion mass tolerance: 10 ppm; MS2 ion mass tolerance: 20 ppm; MS3 ion mass tolerance, 0.6 Da. Maximal number of missed cleavages: 2; minimum peptide length: 6; max. modifications: 4; peptide mass: 350-10000 Da. In the XlinkX node acquisition mode MS2-MS3 was selected and cross-link reporter ion doublets set to Δ-mass 31.9721 Da. Caramidomethylation (+57.021 Da) for cysteins was used as a static modification. DSSO cross-linked mass modifications were used as variable modifications for lysine or protein N-terminus in addition to methionine oxidation (+15.995 Da) and protein N-terminal acetylation (+42.011 Da). Data were searched against a database containing Uniprot/SwissProt entries of the given individual proteins of the eight protein mixture or a reduced human SwissProt database (retrieved Feb. 2019) based on protein identifications from the single-shot proteomics experiment of HEK293T cell lysate. The false discovery rate (FDR) was set to 1% using the Percolator node. Additionally, cross-links were filtered with an identification score of ≥50 and Δ-score of ≥10. Protein abundances were estimated based on iBAQ^22^ and MS1 features were extracted using MaxQuant^23^ 1.6.2a. Search parameters were: MS ion mass tolerance: 10 ppm, MS2 ion mass tolerance: 20 ppm; fixed modification: carbamidomethylation (cystein); variable modification: oxidation (methionine) and acetylation (protein N-term); enzymatic digestion: trypsin; allowed number of missed cleavages: 2; minimum peptide length: 7; database as above; FDR at 1%. All output files were used as input for data analysis in R. The mass spectrometry proteomics data have been deposited to the ProteomeXchange Consortium via the PRIDE^24^ partner repository with the dataset identifier PXD018128.

## Results

### Optimization of FAIMS CV settings using eight cross-linked protein mixture

The FAIMS device enables user-directed fine-tuning mainly at two levels: flow rate of N2 carrier gas and CV. While default settings for the first parameter is suggested by the manufacturer (no adjustment to flow rate), CV values need to be adjusted based on experimental setup. In this study, we first set out to evaluate the CV settings for XL-MS experiments using a medium complex sample. Eight proteins were individually cross-linked, digested and the resulting peptides mixed in equal amounts. The mixture was analyzed by LC/MS with and without FAIMS. Several MS and cross-link characteristics, including the charge distribution of MS1 features, the number of cross-links and the overlap of identifications were evaluated over a range of 10 CVs from -90 V to -40 V, spaced 5 V or 10 V apart.

The charge distribution of MS1 features showed distinctive trends along the selected CV range (Figure 1a and S-2a). The number of MS features at charge states +1 or +2 was the highest at CV -40 and decreased towards lower CV values. In contrast, the number of MS features of charge states +3 maximized at CV -80 and dropped substantially towards higher CV values. Highly charged features (+4 to +7) were enriched in the range of medium to high CVs (i.e., -65 V to -50 V). Furthermore, a strong correlation between sequence length, predicted hydrophobicity and CV was observed in the range of -40 V to -80 V, indicating a selection preference for shorter and more hydrophilic cross-links at lower CVs (Figure S1a, b and S2c, d). Because different CVs select cross-links with different physiochemical properties, we reason that two or three CVs may be used in combination to cover a broader range of cross-linked species.

**Figure 1.**
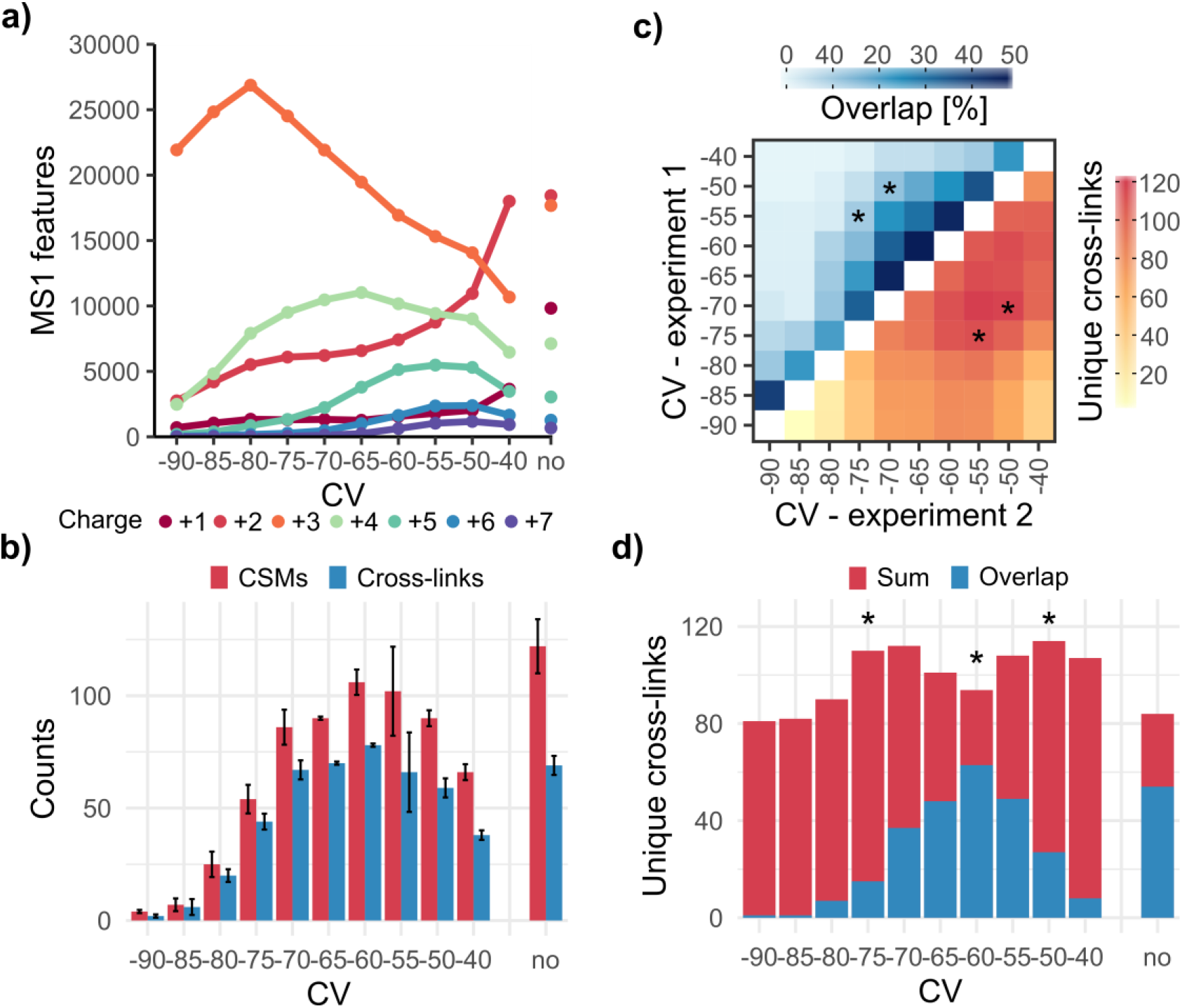
Analysis of FAIMS CV settings using DSSO cross-linked tryptic peptides derived from an eight protein mixture. (a) Number of detected MS1 features along the selected FAIMS CV range. (b) Number of CSMs and unique cross-links for individual CVs. (c) Analysis of sum and overlap of cross-links between two individual CV measurements. * notes the selected 2-CV combinations. (d) Heart plot to determine the best three-CV combination. A first experiment (single CV measurement at -60 V) is combined with a second experiment at different CVs. Local maxima on both sides of the initial CV suggest which three CVs to combine in order to maximize cross-link identification (red) and minimize overlap between CVs (blue). A 3-CV combination (-50/-60/-75 V, marked with *) is selected based on this analysis. Results in (a), (c) and (d) are from replicate 1. Results in (b) are shown as the average of two replicates. In (a), (b) and (d) analysis without FAIMS is indicated with “no”.

First, we analyzed the number of cross-link spectrum matches (CSMs) and unique cross-links of each CV in replicates. Similar to what has been shown in previous proteomic studies applying FAIMS^19,21^, we found that the number of identified cross-links resembled a Gaussian-like distribution across the selected CV range (Figure 1b), reaching a maximum at -60 V. Comparing to the measurement without FAIMS, -60 V yielded a higher number of cross-links while -55 V, -65 V and -75 V provided slightly lower numbers.

Second, we analyzed the total number of cross-links of two individual FAIMS measurements and their overlaps in identification within a CV range of -90 to -40 V in replicates (Figure 1c and S-2b). We selected two 2-CV combinations (i.e., -50/-70 V and -55/-75 V) that provided the best compromise between high numbers of cross-links (118 and 113 cross-links) and low overlaps between CVs (12% and 10% overlaps). 3-CV combinations were surveyed based on similar criteria. In this regard, we selected two CVs (i.e., -55 V and -60 V), which were in the middle of the CV range and provided two high points in cross-link identification, and combined with results from another single-CV measurement ranging from -40 to -90 V. We chose two 3-CV combinations (-40/-55/-70 V and -50/-60/-75 V) by selecting the initial CV value (-55 or -60 V) and the local maxima with the least overlap in identification on both sides (-40 and -70 V for initial CV of -55V; -50 and -75 V for initial CV of -60 V). This method has previously been demonstrated to provide the best estimate for 3-CV combinations.^19^

Third, we analyzed the number of cross-links using internal CV stepping of the selected two 2-CV and two 3-CV combinations. We evaluated three gradient lengths, including 60, 90 and 120 min. We found that the number of CVs to use depends on the gradient length. For 60 gradient, 2-CV combinations resulted in a modest gain of cross-link identification comparing to the measurement without FAIMS. However, this number dropped if 3-CV combinations were used. This is likely due to the saturation of potential cross-link features (higher charged MS1 precursors) when short gradients are applied. For 90 and 120 min gradients, both 2-CV and 3-CV combinations provided better results than the analysis without FAIMS (Figure 2a). Furthermore, these results showed a clear tendency that more CVs can be combined with increasing gradient length to achieve a higher gain in cross-link identification (Figure 2b). For a 120 min analysis the best 3-CV combination (-40/-55/-70 V) increases cross-link identification by 23% (112 without and 138 cross-links with FAIMS).

**Figure 2.**
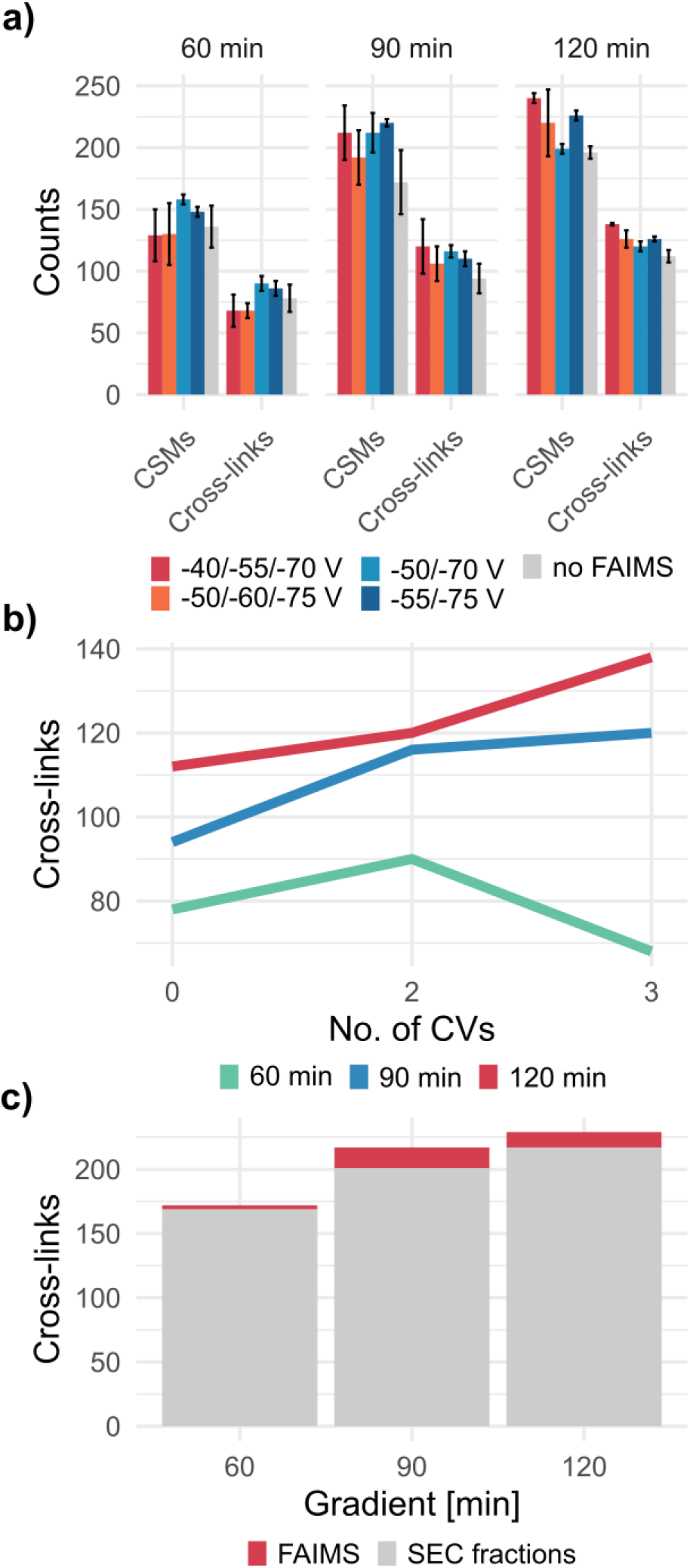
Comparison of different internal CV stepping in combination with different LC/MS gradient lengths. Results are from both replicates of the DSSO cross-linked eight protein mixture. (a) Number of CSMs and cross-links obtained from different CV combinations and LC/MS gradient lengths. (b) Titration of number of CVs over 60, 90 and 120 min gradient lengths. No. of CVs: 0 = no FAIMS, 2 = -50/-70 V, 3 = - 40/-55/-70 V. (c) Comparison of number of cross-links between five SEC fractions (grey) and five different two-CV measurements (-40/-60, -50/-70, -55/-70, -55/-75, -60/-75 V) of unfractionated sample (red). Analyses are performed for 60, 90 and 120 min gradient lengths.

### Comparing cross-link analysis of single-shot FAIMS with SEC fractions

In many previous XL-MS studies of protein complexes, SEC and SCX were used to enrich low abundant cross-links for more efficient cross-link identification.^8,11^ Since FAIMS allows gas-phase separation, we inquire whether FAIMS is capable to substitute chromatography-based separation for medium complex samples. Therefore, we performed SEC to fractionate the peptide digest of the eight cross-linked proteins. In total, we identified 165, 192 and 209 cross-links from five cross-link-enriched SEC fractions using 60, 90 or 120 min gradients without FAIMS. Intriguingly, we were able to achieve a higher number (167, 213 and 223 cross-links) combining five best performing 2-CV FAIMS measurements (-40/-60 V, -50/-70 V, -55/-70, -55/-75 V, -60/-75 V) (Figure 2c). Notably, FAIMS still outperformed SEC fractionation when using a 30 min shorter gradient (90 instead of 120 minutes). This comparison clearly showed that gas phase separation by FAIMS may serve as an easy and effective alternative to in-solution-based fractionation techniques for cross-link identification in medium complex samples.

### Application of FAIMS in proteome-wide XL-MS studies

To evaluate the benefits of FAIMS in highly complex samples, we acquired LC/MS measurements of DSSO-cross-linked cell lysates with and without FAIMS. For this experiment, we prepared lysates from HEK293T cells, cross-linked the cell lysate with DSSO, digested and fractionated with SCX to enrich cross-links. We tested six equally spaced later SCX fractions using the previously optimized two 3-CV combinations (-40/-55/-70 V and -50/-60/-75 V) in a 3-hr gradient. Because of the expected high complexity of the sample, only 3-CV combinations and a longer LC gradient were used, recapitulating a standard LC/MS acquisition for large-scale XL-MS studies.^3,8^

As an overview, we observed a notable improvement by applying FAIMS. The two 3-CV FAIMS measurements yielded 3,491 and 3,484 cross-links, providing a 56% increase compared to the 2,231 cross-links obtained from the same analysis without FAIMS (Figure 3a). This improvement was observed throughout all six SCX fractions, however was the greatest for the first two fractions (more than two fold), and dropped to ~40% for the later four fractions (Figure 3b, c). This is likely due to the decrease of sample complexity along the SCX gradient. To further dissect the contributions of each CV in the measurements, we analyzed cross-link identification of each CV and the overlaps between CVs. We found that CVs around the initially observed identification maximum in single-CV measurements (i.e., -50, -55, -60 and -70 V) contributed to more than 60% of all identified CSMs, while the third more distant CV (-40, and -75 V) contributed to less than 30% (Figure S-3a). In addition, we noted that a spacing of 15 V reduced the overlap of identifications between CVs compared to a spacing of 10 V (Figure S-3b, c).

**Figure 3.**
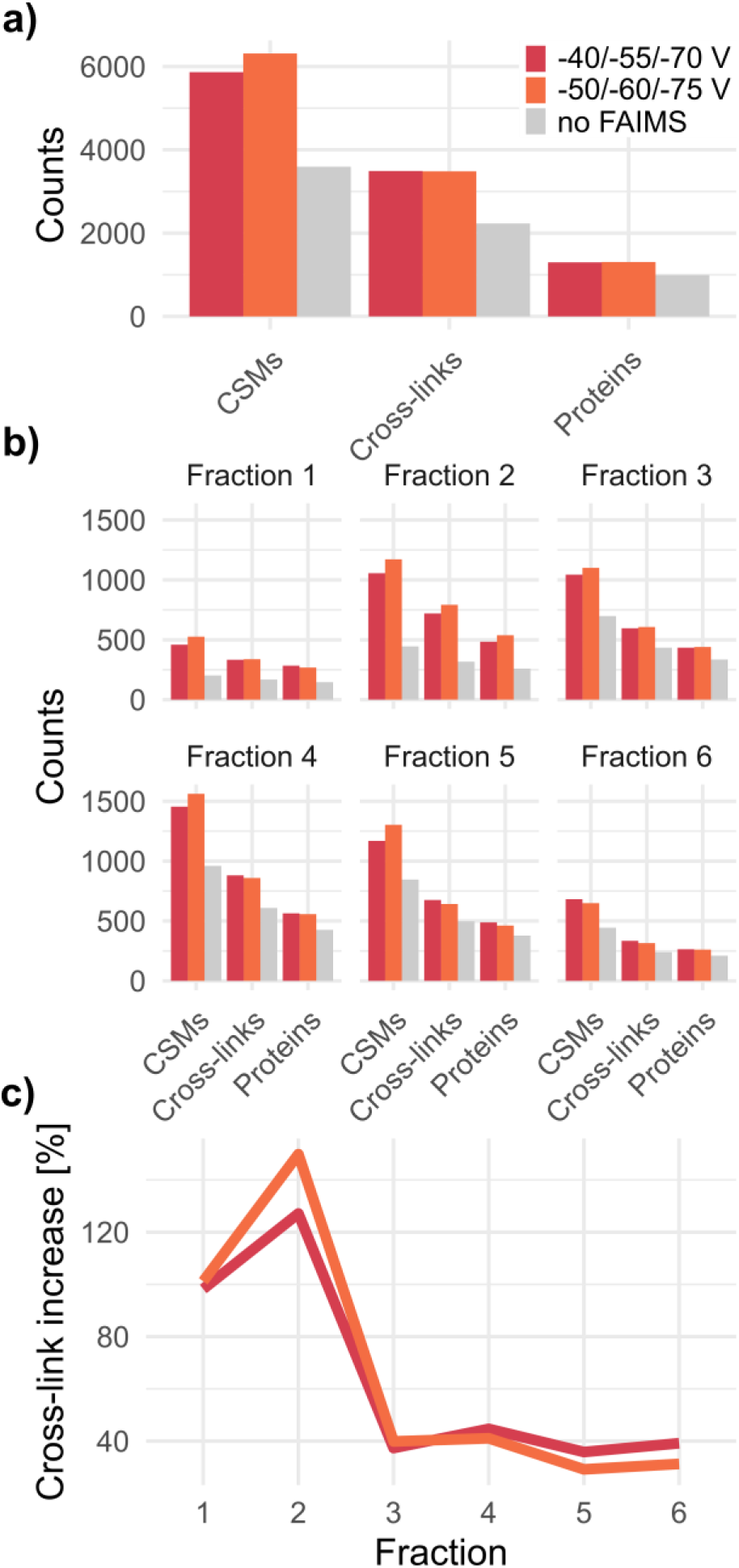
Characterization of FAIMS using six SCX fractions from DSSO cross-linked HEK293T cell lysate. (a) Number of CSMs, unique cross-links, and cross-linked proteins over six SCX fractions using two 3-CV FAIMS settings and without FAIMS. (b) Results in a) shown per individual SCX fraction. (c) Percentage increase of cross-link identification in different SCX fractions using FAIMS.

Next, to analyze the contributing factors of this improvement, we assessed several important MS characteristics and quality measures. First, we analyzed the number of MS1 features and the charge distributions across the six selected SCX fractions. In all FAIMS settings, features at charge state +1 and +2 (depending on CV also +3) were greatly suppressed, while the amount of higher charged features relative to all MS1 features increased substantially (Figure 4a). This result clearly showed that FAIMS led to a relative enrichment of highly charged features over undesired lower charged ones in cross-link analysis. Second, we assessed three important spectral quality indicators, including cross-link identification rate, MS1 isolation interference and cross-link identification score (XlinkX score). Our analysis showed that FAIMS led to a higher cross-link identification rate for all fractions (Figure 4b). It also significantly improved MS1 isolation interference for all fractions and XlinkX score for early SCX fractions (Figure 4c, d), suggesting better MS2 spectral quality when applying FAIMS.

**Figure 4.**
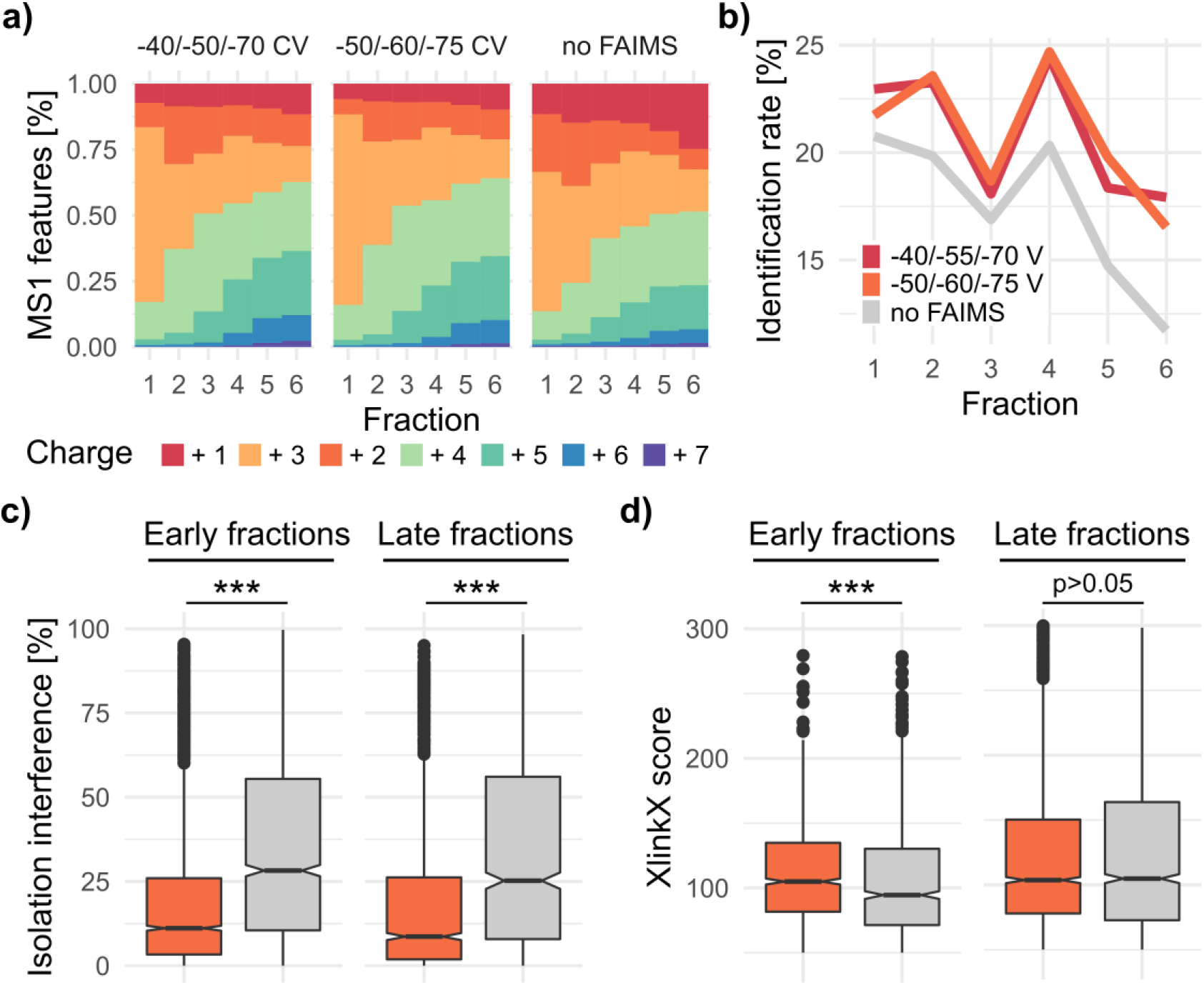
Assessment of MS parameters and quality measures. (a) Distribution of MS1 charge states for the two selected 3-CV combinations. (b) Analysis of identification rate (percentage of cross-link identifications from all sequencing events) in different SCX fractions and FAIMS measurements. (c) Distribution of MS1 isolation interference and (d) cross-link identification score (XlinkX score) for the best-performing FAIMS setup (-50/-60/-75 V, orange) and without FAIMS (gray). Only cross-links identified in both measurements are selected for the analysis. Early fractions: fractions 1-3; late fractions: fractions 4-6. Significance is calculated by two-sided Wilcoxon rank-sum; ***p < 0.0001.

In addition to the advancements at cross-link level, we ask how FAIMS improves detection of cross-linked proteins and depth of the interactome. At the level of protein-protein interaction, we observed a 50% increase in intra-links, 70% increase in inter-links and 56% increase in cross-linked protein pairs (Figure 5a, b). These numbers were highly similar between the two tested 3-CV combinations. Importantly, FAIMS expanded the detection of lower abundant proteins to a larger extend compared to higher abundant ones (Figure 5c, d), thus emphasizing its added value in detection of low abundant protein-protein interactions in large-scale XL-MS studies.

**Figure 5.**
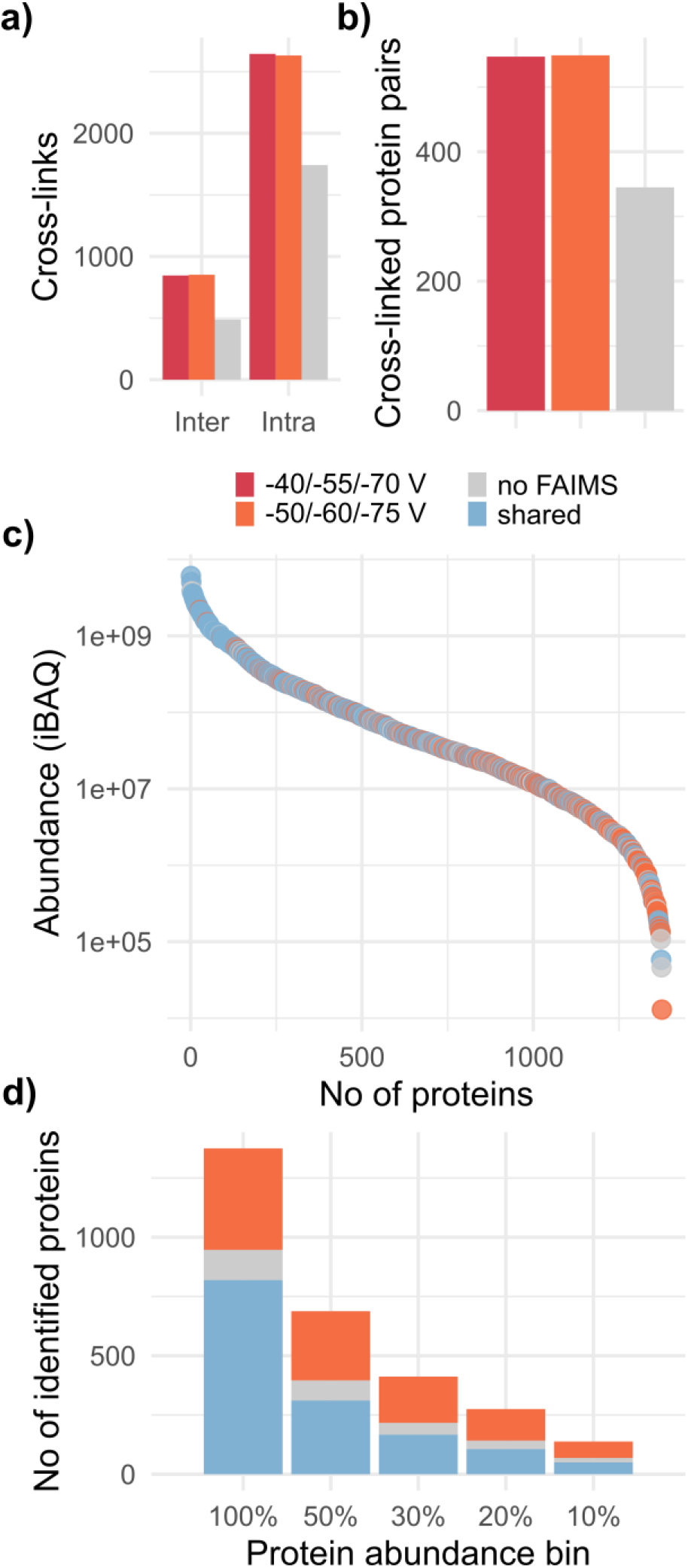
Assessment of FAIMS at the level of protein-protein interactions. (a) Total number of inter- and intra-links from the six selected SCX fractions in different (FAIMS) measurements. (b) Total number of unique cross-linked protein pairs. (c) Cross-linked proteins ranked by iBAQ values and color coded by identification method. Orange: unique identifications from -50/-60/-75 V measurement; grey: no FAIMS; blue: cross-links identified in both FAIMS analysis. (d) Number of cross-linked proteins in different abundance bins based on their iBAQ values. 100% corresponds to all and 10% to the lowest abundant proteins. Colors are the same as in (c).

## Discussion

Over the past decade, XL-MS has been widely used as a low-resolution structural technique to probe the architecture of multi-protein complexes in combination with other means in structural biology. In recent years, it gained increasingly interest in large-scale interactome studies to profile protein-protein interaction networks and discover novel protein partnerships. Therefore we evaluated the performance of FAIMS in cross-link identification based on these two types of samples. We prepared a medium complex sample by mixing eight individually cross-linked proteins and a high complex sample from SCX-fractionated cell lysates. We showed that FAIMS increased cross-link identification by 23% in the cross-linked eight protein mixture and 56% in the SCX fractionated cell lysates. This improvement was contributed by several levels of MS measures, such as MS1 isolation interference, cross-link score and identification rate. Furthermore, we evaluated different CV settings in both types of cross-links samples. In agreement with previous observations for linear peptide identification, we found that the ideal number of CVs to use depends on the gradient length. One CV per hour seems to be a reasonable estimation. For complex samples, an internal 3-CV stepping of -50/-60/-75 V or -40/-55/-70 provided similarly good results in a three-hour gradient.

Importantly, our results on medium complex samples showed that FAIMS boosted the number of cross-link identification to a similar level as compared to that using SEC fractionation, even with shorter LC gradients. Single-shot FAIMS measurement requires much lower sample amount, reduces sample preparation time and MS instrument time. Therefore it is highly beneficial to XL-MS studies of protein complexes especially in cases where the amount of proteins are limited.

For complex samples, in addition to the global improvements as mentioned above, we questioned whether SCX fractions at different elution times affect optimal CV settings. In this respect, we evaluated several physiochemical properties of cross-links in the SCX fractions. As expected, we observed a decrease of hydrophobicity and sequence length as well as an increase of charge states along the SCX gradient (Figure S-4a-c). However, these differences in physiochemical properties of cross-links did not offer additional benefit in gas-phase separation by FAIMS. Only minor differences in cross-link identification were observed between the two 3-CV combinations across all six SCX fractions (i.e, slight preferences for -50/-60/-75 V in the first three SCX fractions and -40/-55/-70 V in the last three SCX fractions) (Figure 3b). Using a 3-CV combination adjusted to more hydrophilic and shorter peptides (-60/-70/-80 V) we did not observe any benefits along the SCX fractions (Figure S-5a, b). Furthermore, we generated cross-link density maps along the LC gradient for each SCX fraction to evaluate the contribution of each CV at different LC retention time (Figure S-4d, e). Interestingly, we observed a notable shift of cross-link density to the frontend of the LC gradient in the first two SCX fractions. A possible explanation is that the early SCX fractions contain a higher percentage of linear peptides that are more susceptible to separation from cross-links by FAIMS. For the other SCX fractions, the cross-link density maps are mostly similar to the ones without FAIMS. These analyses indicated that with the current FAIMS setup, the resolution may not be sufficient to further separate the cross-links based on their physiochemical properties. However, a higher resolution FAIMS device will likely further improve the number of cross-link identification by fine-tuning the CV combinations in different SCX fractions as well as at different retention time of the LC gradient.

## Supporting Information

Fig. S1. Additional analysis of DSSO cross-linked eight protein mixture (replicate 1).

Fig. S2. Analysis of DSSO cross-linked eight protein mixture (replicate 2).

Fig. S3. Analysis of cross-link identifications per FAIMS CV using six SCX fractions from DSSO cross-linked HEK293T cell lysate.

Fig. S4. Analysis of physiochemical properties of cross-links along SCX fractions and LC/MS retention time using six SCX fractions from DSSO cross-linked HEK293T cell lysate.

Fig. S5. Characterization of one additional FAIMS setup using six SCX fractions from DSSO cross-linked HEK293T cell lysate.

## Author Contributions

L.S. performed the experiments and data analysis. S.S performed the SEC experiment. M. N., C. F. and R. V. assisted data acquisition. L.S., and F.L. wrote the manuscript; F.L. supervised the research. All authors reviewed the manuscript.

## Notes

RV is employee of Thermo Fisher Scientific.

The authors declare no competing financial interest.

## Acknowledgements

We gratefully acknowledge support from Deutsche Forschungsgemeinschaft Grant SFB958. We thank Thermo Fisher Scientific, and especially Michael Belford, and Max Planck Unit for the Science of Pathogens for the access and support of the FAIMS Pro™ device coupled to the Orbitrap Fusion Lumos mass spectrometer.

## Insert Table of Contents artwork here

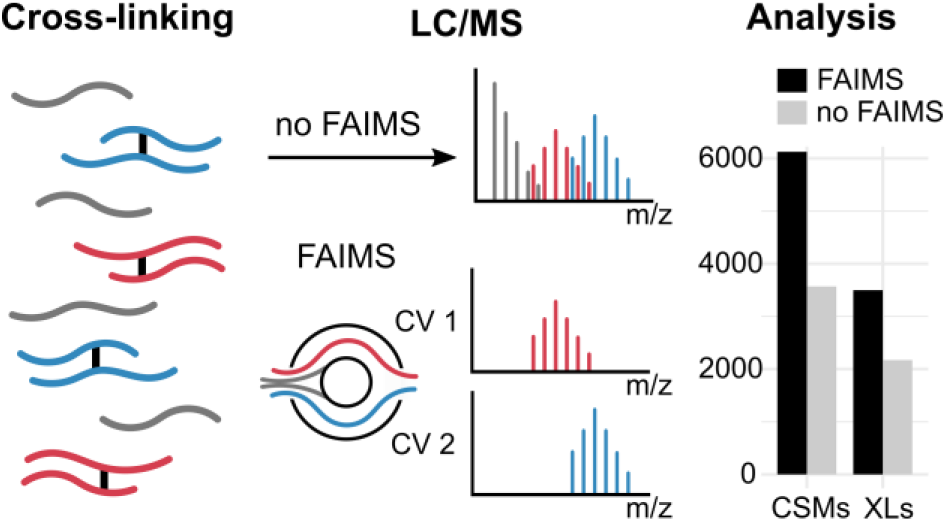

