## Supporting Information and Figures for "Expanding the depth and sensitivity of cross-link identification by differential ion mobility using FAIMS"

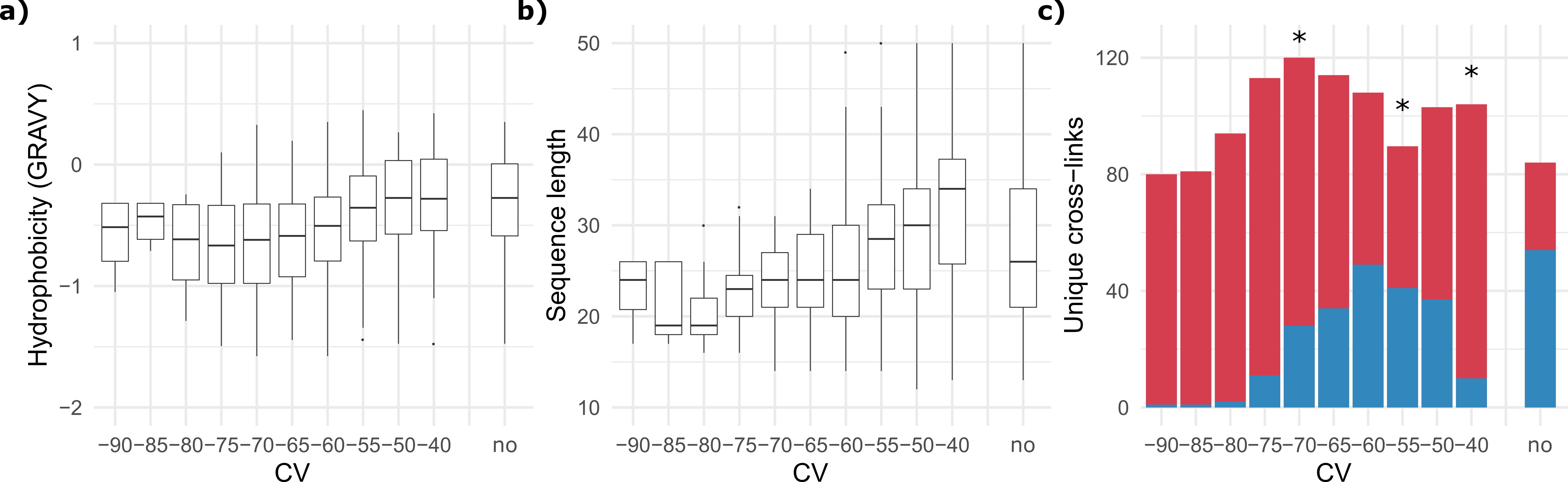

**Supplementary Figure 1.** Additional analysis of DSSO cross-linked eight protein mixture (replicate 1). (a) Distribution of predicted hydrophobicity based on Gravy index score (> 0 hydrophobic, < 0 hydrophilic) along selected CV range. (b) Distribution of the length of linearized cross-link sequence along the selected CV range. (c) Heart plot to determine the best three-CV combination. A first experiment (single CV measurement at -55 V) is combined with a second experiment at different CVs. Local maxima on both sides of the initial CV suggest which three CVs to combine in order to maximize cross-link identification (red) and minimize overlap between CVs (blue). A 3-CV combination (-40/-55/-70 V, marked with *) is selected based on this analysis. Analysis without FAIMS is indicated with “no”.

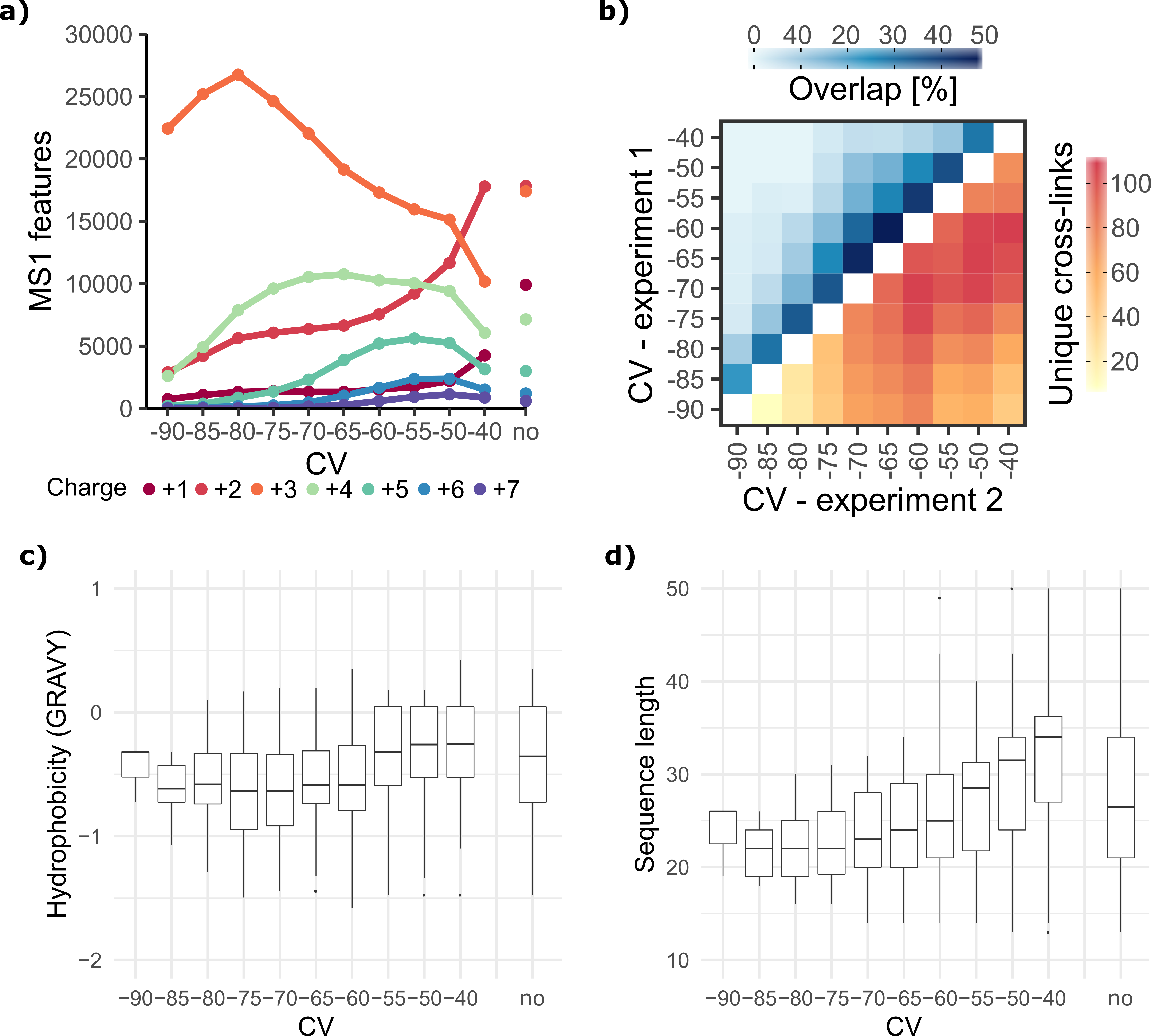

**Supplementary Figure 2.** Analysis of DSSO cross-linked eight protein mixture (replicate 2). (a) Number of detected MS1 features along the selected FAIMS CV range. (b) Analysis of sum and overlap of cross-links between two individual CV measurements. (c) Distribution of predicted hydrophobicity based on Gravy index score (> 0 hydrophobic, < 0 hydrophilic) along selected CV range. (d) Distribution of the length of linearized cross-link sequence along the selected CV range.

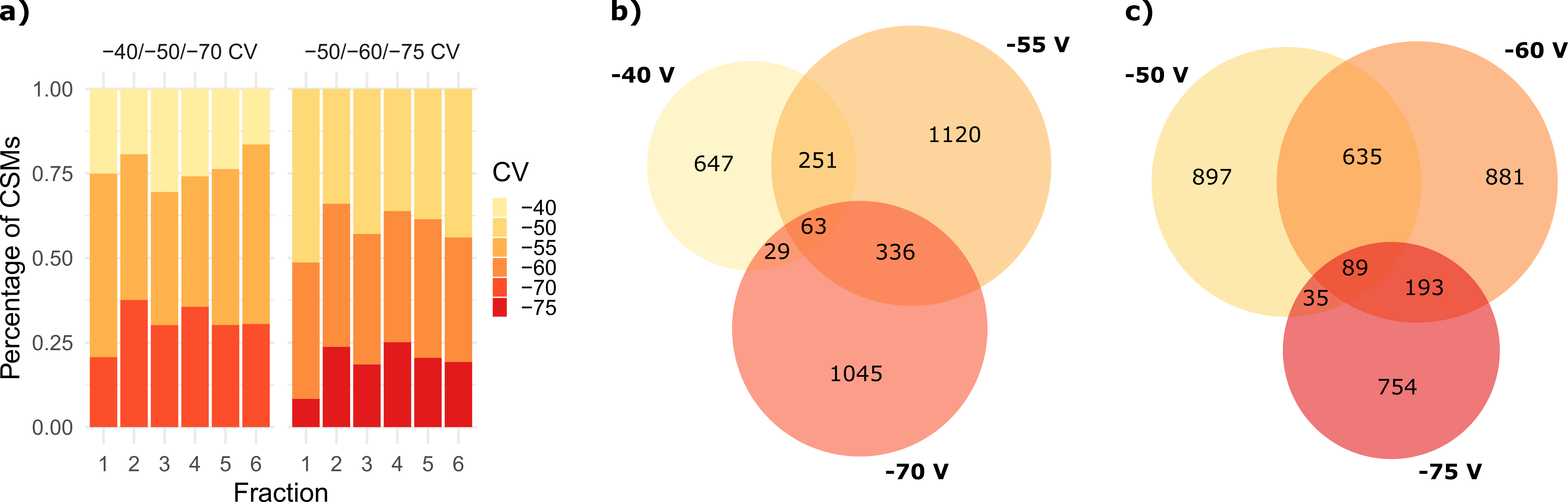

**Supplementary Figure 3.** Analysis of cross-link identification of individual FAIMS CV using six SCX fractions from DSSO cross-linked HEK293T cell lysate. (a) Percentage of CSMs identified by each CV within one FAIMS internal stepping along SCX fractions. (b, c) Overlap of cross-links between different CVs in (b) -40/-55/-70 V and (c) -50/-60/-75 V measurements over all SCX fractions.

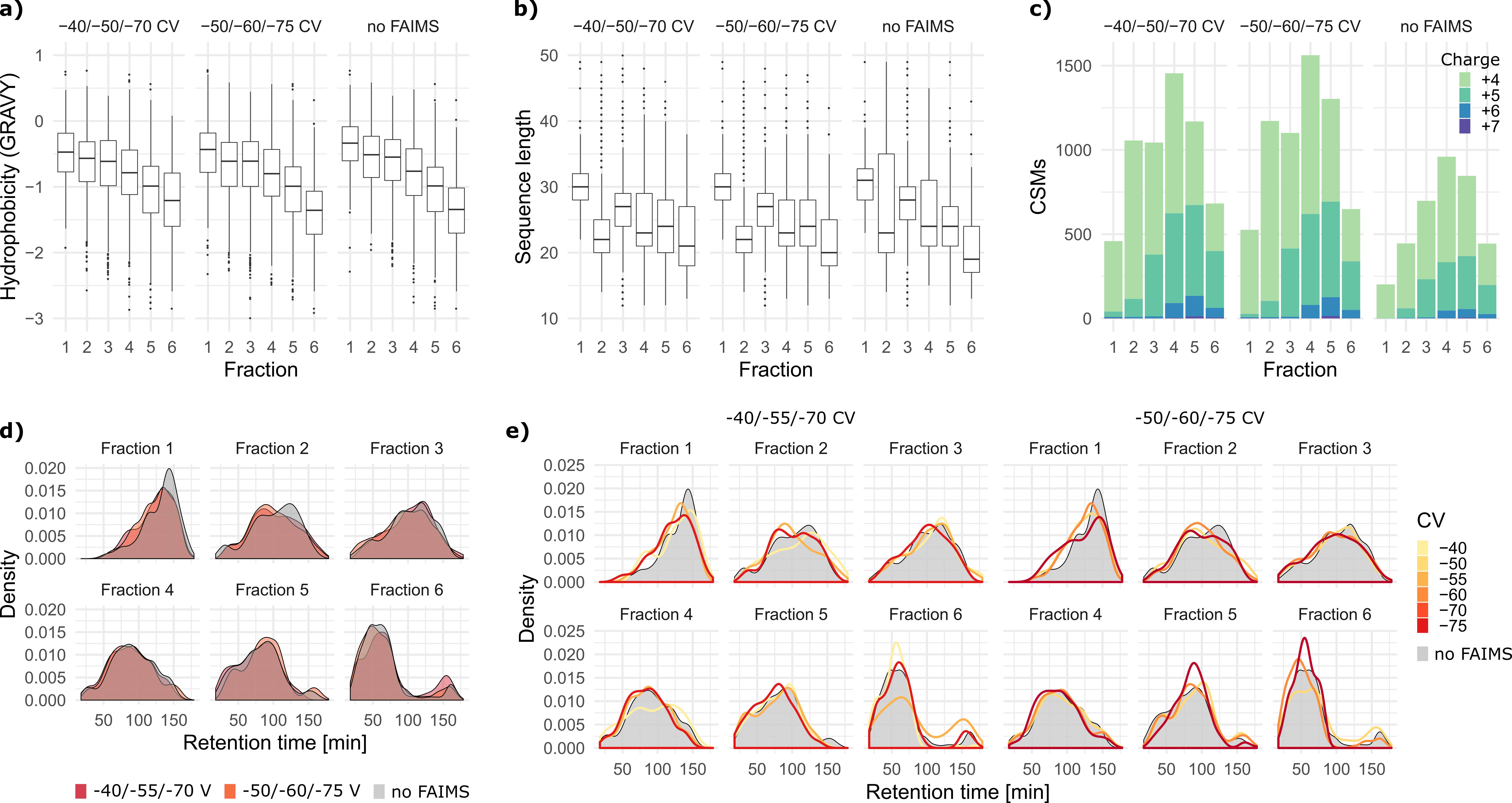

**Supplementary Figure 4.** Analysis of physiochemical properties of cross-links along SCX fractions and LC/MS retention time using six SCX fractions from DSSO cross-linked HEK293T cell lysate. (a, b) Distribution of (a) predicted hydrophobicity based on Gravy index score (> 0 hydrophobic, < 0 hydrophilic) and (b) distribution of the length of linearized cross-link sequence along SCX fractions in different FAIMS setup. (c) Number of CSMs in different charge states along SCX fractions in different FAIMS setup. (d, e) Density distribution of CSMs along LC/MS gradient. Results of 3-CV combinations are shown in (d) and individual CVs are shown in (e).

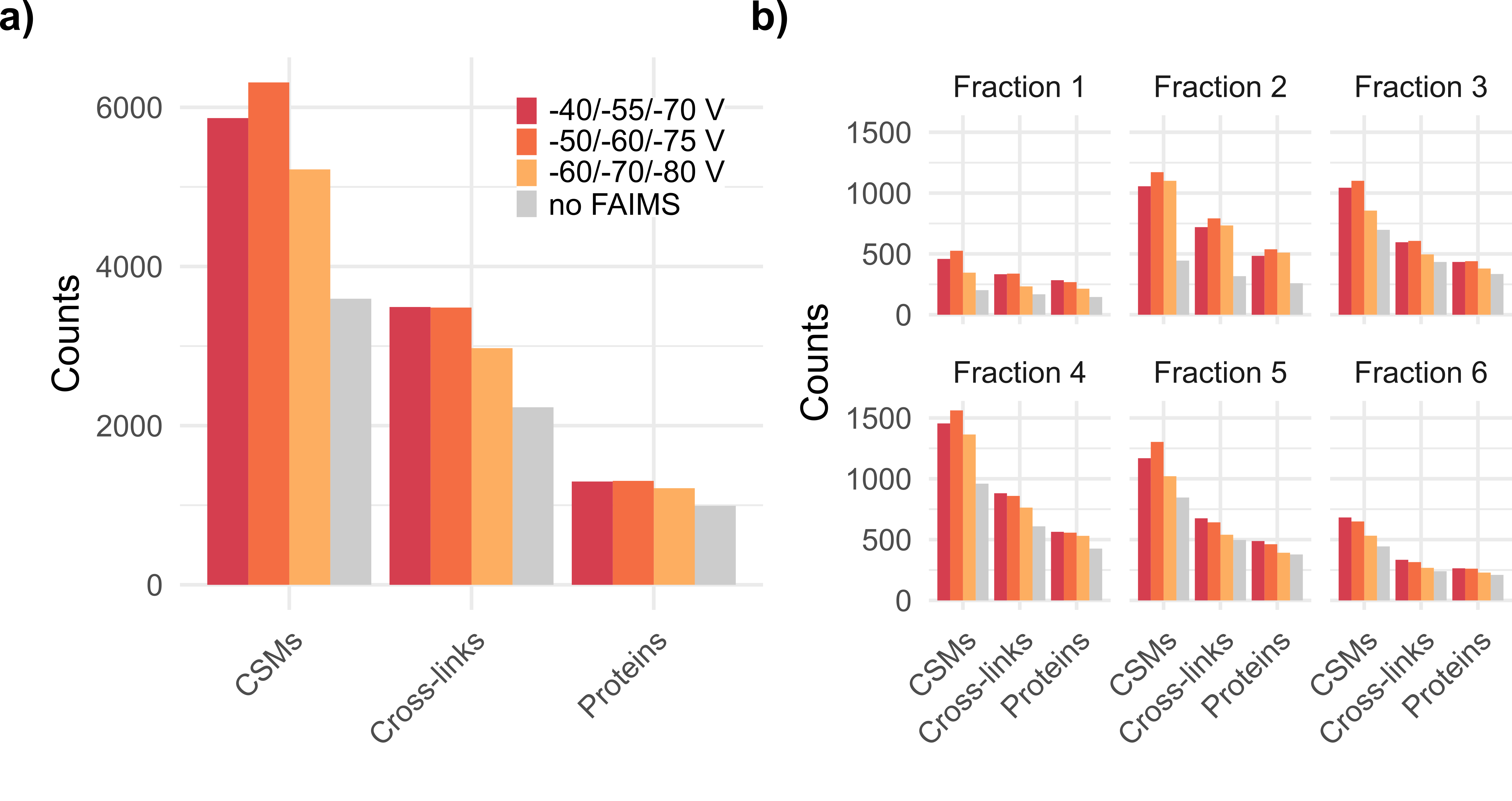

**Supplementary Figure 5.** Characterization of one additional 3-CV combination using six SCX fractions from DSSO cross-linked HEK293T cell lysate. (a) Number of CSMs, unique cross-links, and cross-linked proteins over six SCX fractions using three 3-CV FAIMS settings and without FAIMS. (b) Results in a) shown in individual SCX fraction.
